# Hound: A novel tool for automated mapping of genotype to phenotype in bacterial genomes assembled de novo

**DOI:** 10.1101/2023.09.15.557405

**Authors:** Carlos Reding, Naphat Satapoomin, Matthew B. Avison

## Abstract

Increasing evidence suggests that microbial species have a strong within species genetic heterogeneity. This can be problematic for the analysis of prokaryote genomes, which commonly relies on a reference genome to guide the assembly process. Any difference between reference and sample genomes can introduce errors in the detection of small insertions, deletions, structural variations and even point mutations. This phenomenon jeopardises the genomic surveillance of antibiotic-resistant bacteria, with predictions of resistance varying between laboratories. Here we present Hound, an analysis pipeline that integrates publicly available tools to locally assemble prokaryote genomes *de novo*, detect genes by similarity using the proteins they encode as query, and report the mutations found. Three features are exclusive to Hound: it reports relative gene copy number, retrieves sequences upstream the start codon to detect mutations in promoter regions—which allow gene expression signals to be integrated—and, importantly, can merge contigs based on a user-given query sequence to reconstruct genes that are fragmented by the assembler. To demonstrate Hound, we screened through 5,032 bacterial whole-genome sequences isolated from farmed animals and human infections, using the amino acid sequence encoded by *bla*_*TEM-1*_, to predict resistance to amoxicillin/clavulanate which is driven by over-expression of this gene. We believe this tool can facilitate the analysis of prokaryote species that currently lack a reference genome, and can be scaled up to build automated systems for antibiotic susceptibility prediction.

## I. INTRODUCTION

The advent of affordable genome sequencing has exposed the wide genetic heterogeneity that exists within bacterial species [1]. With genome sizes that range between 2.69–2.92 Mb in *Staphylococcus aureus*, or between 4.66–5.30 Mb for *Escherichia coli*, it is not surprising that some begin to question the notion of *species* [2, 3] or even *clone* [4] in prokaryotes. This heterogeneity led to the concept of *pan-genomes* [5], but it also exposes another, more technical problem: How to study the genomes of prokaryotes without masking this genetic diversity?

Raw sequencing data are typically mapped onto a high-quality reference—whose sequence is known and resolved (i.e. circularised) [6, 7]—or databases containing them [8], to study the genetics of organisms from viruses [9] to vertebrates [10] or plants [11]. The use of reference-mapped assemblies is used in comparative genomics [12], clinical microbiology [13], public health [14, 15], and even to inform policy through the detection of specific mutations or phylogenetic analyses [16, 17]. Now, given the further reduction in sequencing costs, reference-mapped assemblies are increasingly used to predict antibiotic susceptibility in bacteria from clinical samples [18, 19]. This is driven by the ability of genomics to screen resistance to multiple antibiotics simultaneously, more than is possible with current phenotypic antibiotic sensitivity tests, improving antibiotic stewardship and patient care. But using genomics data for this can be problematic, given the limitations of these type of assemblies to detect antibiotic-resistance genes. On one hand, reads that cannot be mapped onto the reference genome, say, because they are plasmid-borne and not part of the chromosome, are excluded from the assembly. And this loss of data hinders the detection of antibiotic-resistance genes [19, 20]. On the other hand, the availability of reference genomes is skewed towards the most common pathogens [13, 16], further limiting the study of rarer pathogens [21]. Consequently, the scope of tools like ResFinder [22], STARR [23], ARG-ANNOT [24], RAST [25], or ABRIcate [26] can be limited to predict antibiotic susceptibility. Particularly, because they rely on reference genomes to report the presence—or not—of antibiotic-resistance genes along with mutations in the coding sequence *known* to be associated with specific resistant phenotypes. As we show below, mapping sequencing data onto a reference genome can artificially modify the assembly [20]. This approach is not only limited for the study of other pathogens or the finding of novel, undocumented mutations associated with important phenotypes like antibiotic resistance; but of species that may have other biological or ecological importance where reference genomes and tools are scarce [27].

Here we sought to build a pipeline to analyse bacterial genomes assembled *de novo*, without using a reference to guide the assembly process. *De novo* assemblies lack most of the limitations mentioned above, but can also introduce others. Particularly, the fragmentation of genes—whose sequences are split across multiple contigs by the assembler [19]. Hound implements an algorithm to re-purpose a query sequence as a local reference to detect and merge the relevant contigs, so that its sequence can be reconstructed unambiguously. Another issue we sought to address is that antibiotic resistance is not only caused by the presence of specific genes. The over-expression of antibiotic resistance genes, whether through specific mutations in the promoter or increase in relative gene copy number, can dramatically alter the antibiotic resistance phenotype present. For example, amoxicillin-resistant *Escherichia coli* are most commonly resistant due to the production the TEM β−lactamase enzyme, encoded by the mobile gene *bla*_*TEM-1*_ [28]. Amoxicillin given in combination with clavulanate will kill amoxicillin-resistant *E. coli* because clavulanate inhibits TEM-1—as well as other related enzymes—explaining why this combination has been widely used in human [29, 30] and veterinary medicine [31]. However, *E. coli* can become resistant to amoxicillin-clavulanate by over-producing TEM-1 due to promoter mutations [29] or increased gene copy number [32]. Therefore, we built into Hound the capability to retrieve sequences beyond a gene’s coding sequence and include the promoter, as well as the relative gene copy number, to allow the detection of such variants with our pipeline.

## II. RESULTS

### Pipeline overview

Hound integrates tools widely-used to assemble Nanopore and Illumina reads *de novo*, and screen the resulting assemblies for user-given query sequences, into a single tool. Hound supports nucleotide and amino acid sequences, but we suggest the query to be an amino acid sequence where possible to avoid variations introduced by synonymous mutations. Our pipeline is modular as Figure 1 illustrates to allow performing only a subset of the tasks, and relies on SPAdes [33] as its backend assembler due to its combination of speed, accuracy, and support for sequencing data from multiple platforms. Once assembled, Hound can optionally map the raw reads onto the assembly using Burrows-Wheeler aligner [34] to compute the coverage depth with SAMtools [35]. Following the assembly step Hound will search for the query sequences in the assembly by similarity, using the BLAST [36] algorithm, before undergoing downstream processing. At this point, Hound will integrate the data to estimate the relative gene copy number, retrieve sequences upstream of the coding sequence in the assembly, align and produce a phylogeny of all retrieved sequences, and detail the mutations with respect the query sequence.

**Figure 1.**
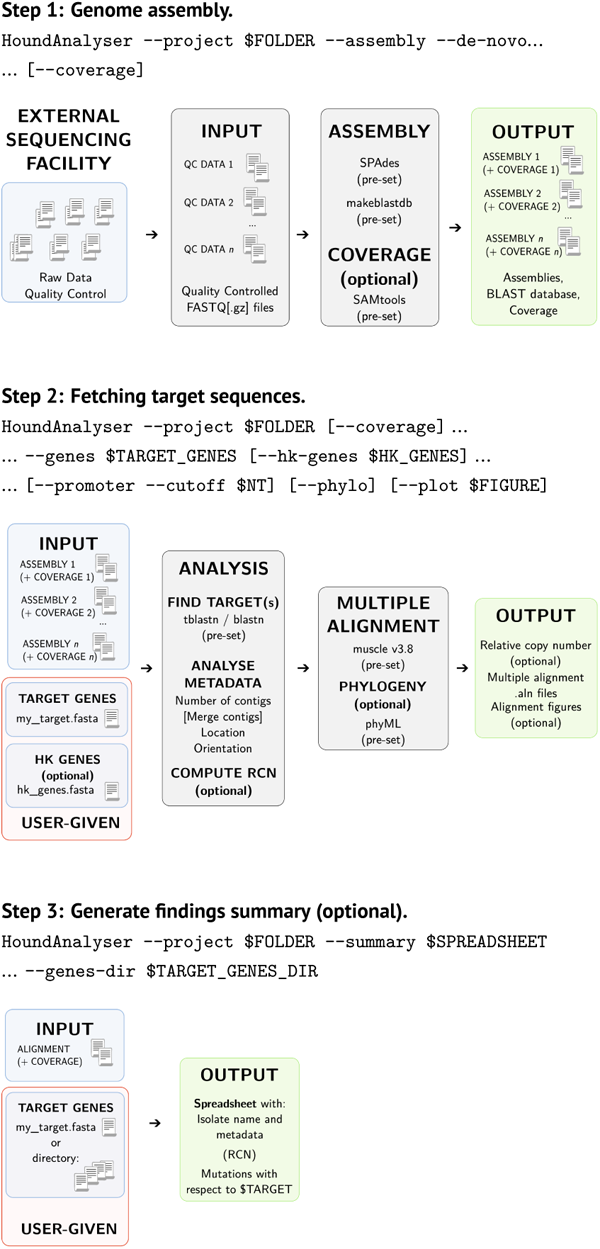
Description of the Hound pipeline for the analysis of bacterial genomes assembled *de novo*. While Hound can be run in a single step, the user is given three steps of granularity. **1)** The first step is to assemble the quality-filtered FASTQ files and depth of coverage data generated. During this step, each assembly is converted into a BLAST database to facilitate downstream analyses. Note that once assembled, there is no need to repeat the step—whence the choice of granularity. **2)** In the second step, Hound will search by similarity any user-given target(s) in the assemblies generated in 1). The target sequence(s) must be in FASTA format and preferably be the amino acid sequence to avoid variations introduced by synonymous mutations. Files with multiple entries are supported. To compute the relative gene copy number (RCN), Hound will use a number of house-keeping genes to compute a baseline depth of coverage. We used four but there is no limit in number of house-keeping genes. If the identity of the sequence found in the assembly with respect to the target, translated as necessary, is above 90%, and the sequence is fragmented, Hound will use the user-given sequence as a guide to sort, de-duplicate, and merge the relevant contigs so the resulting translation is the query sequence used. All sequences are then aligned to facilitate the screening of mutations, insertions or deletions with respect to the target sequence. **3)** The last step is optional, and invokes Hound to summarise all the findings from 2) into one spreadsheet for record-keeping.

The main drawback of *de novo* assemblies [19] is the fragmentation of genes, with subsets of their sequence being split across two or more contigs by the assembler. In this case, if the identity of the sequences found by Hound are *at least* 90% (`$MIN_ID` ≥ `0.9`) to the user-given query, and the contigs harbouring subsets of the query have overlapping common sequences, Hound will shortlist the contigs for its contig-merging routine. Here, after sorting them first by identity and length, Hound will iteratively run pairwise alignments between the first pair with overlapping coordinates, discard one of the two overlapping sequences to avoid introducing duplications, and compare the resulting sequence to the user-given query. Hound will process the next contig in the list and add it to the previous merged sequence until the reconstructed sequence has, at least, equal length to the query which will have at least `$MIN_ID` identity. Note all these contigs will have overlapping coordinates that are located at the boundaries of the contigs, thus, only genes truncated by the assembler and not by insertion sequences—mobile elements—will be included in this analysis.

At this point, once the query sequence can be reconstructed, the assembly is rewritten with the new contig name being a concatenation of all the founding contigs (i.e. `>NODES_1+43+24`). If coverage data exists, the coordinates used earlier by the merging routine are applied to this dataset to preserve coverage in the new, merged contig.

### Reference-mapped *vs de novo* assemblies

The first step in screening recently-acquired genomes of our 5,032 bacterial isolates from farmed animals and human clinical samples, is the assembly *de novo* of the reads. A first look at the output of the pipeline reveals the assemblies seldom have the same size (Figure 2A). Now, while *E. coli* is by far the most abundant species in our dataset, as identified by Kraken 2 [37], the dataset also contains isolates of *Klebsiella pneumoniae* (15.7% of the total), and one isolate of *Pseudomonas aeruginosa* (≪1%) and *Salmonella enterica* (≪1%) among the human clinical isolates. This means multiple species can be monitored depending on each use case.

**Figure 2.**
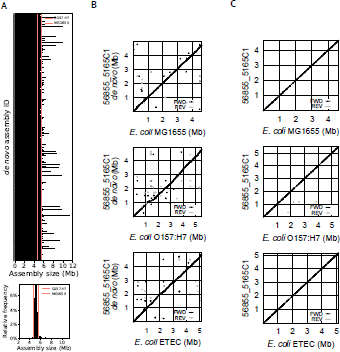
Variability of within-species genome size is not captured by the canonical use of reference genomes. **A)** Histogram (top) and distribution (bottom) of genome sizes across 3562 farm and clinical *E. coli* isolates. The canonical size of the reference wild-type (K-12 substr. MG1655, genome `GCF_000005845.2` in the NCBI) and Shiga toxin-producing strain (0157:H7 str. Sakai, genome `GCA_000008865.2` in the NCBI) are marked at approx. 4.64 Mb and 5.59 Mb in red-toned vertical lines. **B– C)** Dotplots of one representative isolate to visualise the genome-genome sequence alignment between *de novo* (B) or reference-mapped (C) assemblies and three different known genomes: MG1655, O157:H7, and enterotoxigenic *E. coli* (ETEC). Sequences present in these references, but not in out isolate, are noted by a horizontal gap in the dotplot, whereas the converse is noted by a vertical gap. Note that in C) the reported assembly size of our isolate, in the *y* −axis, varies depending on the choice of reference—size is constant in those genomes assembled *de novo*.

When we removed all non-*E. coli* from the dataset and compared the assembly sizes, we found the variation with respect to the available reference genomes—K-12 MG1655 and 0157:h7 Sakai—to be substantial as the histogram in Figure 2A shows, with most assemblies having sizes in-between these references. These two genomes are the only ones validated by the National Center for Biotechnology Information (NCBI) and flagged as reference genomes accordingly. The variability that we observed in genome sizes is consistent with the aforementioned notion of pan-genomes. Next, we compared reference-mapped assemblies using to different *E. coli* genomes using `--assemble --reference $REF_GENOME` with their respective reference—beyond the above MG1655 and 0157:h7 genomes we also used that for enterotoxigenic *E. coli* (ETEC)—as well as the *de novo* assemblies, using pairwise alignments with MUMmer [38].

When comparing the *de novo* assemblies to the references, this alignment revealed signatures of small deletions, insertions, and repeated regions that were different with each reference used (Figure 2B). However, the signatures vanished when we used reference-mapped assemblies (Figure 2C). This means the use of references discards or includes details from the final assembly that can ultimately alter its size—unique and robust when assembled *de novo*—and help explain the inconsistent results [39] that occur when comparing different tools.

### Monitoring *bla*_*TEM-1*_ in farm and clinical isolates

We used this tool to screen through our 5,032 isolate whole-genome sequencing (WGS) datasets, from farmed animals (2,494) and human clinical samples (2,538), to detect *bla*_TEM-1_ (including non-synonymous variants), its promoter region, and relative copy number. Among the output files generated by Hound, there is a figure with the multiple alignment and phylogeny of the sequences, containing the aforementioned metadata depending upon the flags provided, that can be produced with the flag `--plot $FILENAME`. This is an exemplar of how this tool can improve the prediction and, therefore, the surveillance of antibiotic resistance with complex phenotypes that are notoriously difficult to predict from mutations in the coding sequence alone.

The result shows 39.16% (n=994) of clinical isolates were *bla*_*TEM-1*_ positive compared to 51.84% (n=1,283) in those from farm animals (Figures 3B and C). It is noteworthy to mention that human clinical isolates largely came from routine surveillance of Gram-negative bacteria and include multiple species beyond *E. coli* that less commonly carry *bla*_*TEM-1*_, and would also include isolates resistant to very few antibiotics, whereas those from farmed animals were *E. coli* isolates sequenced due to their resistant phenotype (i.e. those recently reported by [40]). Interestingly however, only 22.52% (n=289) of the isolates from farmed animals harboured two or more copies of the gene. This contrast with data from the clinical isolates, where 39.33% of the *bla*_*TEM-1*_ positive isolates (n=391) harboured two or more copies. Since increased gene copy number is associated with amoxicillin/clavulanate resistance [32], this would fit with a higher rate of resistance to this combination given its more widespread use in the clinic to treat humans. Moreover, as Figures 2A and B illustrate, in some isolates *bla*_*TEM-1*_ number is in the hundreds. While their occurrence is rare—n=6 in farm isolates, n=31 in clinical isolates—it suggests the circulation of one or more *bla*_*TEM-1*_-encoding plasmids with very high copy number. Indeed, multicopy plasmids with hundreds of copies are not unheard of [41]. Using the flag `--promoter --cutoff 250` we used Hound to retrieve the promoter of *bla*_*TEM-1*_ and flag any mutation found. Again, mutations associated with over-expression, annotated in the figure produced as black dots, were more common in isolates from humans than farmed animals as Figures 3B and C illustrate. Mutations found by Hound with respect to the query sequence used, as well as relative copy number and other metadata can be exported into a spreadsheet by using the flag `--summary $SPREADSHEET`, which can then be parsed to highlight isolates with specific mutations. An illustrative spreadsheet is included as a supplementary table.

**Figure 3.**
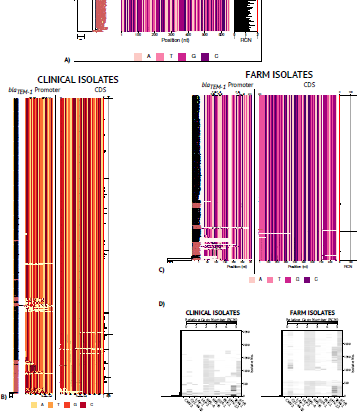
Screening *bla*_*TEM-1*_ across thousands of bacterial isolates. **A)** Summary plots generated with Hound showing phylogeny (left), multiple alignment of the promoter and coding sequence (centre), and relative copy number (RCN, right) of clinical (**B**) and farm isolates (**C**) that are positive for *bla*_*TEM-1*_. A black dot is placed on positions where the nucleotide sequence deviates from the consensus sequence (present in at least 80% of the isolates). Note these are mutations within the multiple alignment, the mutations reported in Hound’s spreadsheet are those from pairwise alignments between the sequence in each isolate and the user-given target sequence and the resulting change in amino acid. Isolates with mutations are highlighted in red by default. When regions of interest have been given alongside the `--promoter` flag, they are added at the top of the alignment (here are promoter regions P_a_-35, P_b_-10, P_3_-35, P_3/4_-10), and start codon. **D)** Heatmap created *ad hoc* to summarise the output of Hound when used to find multiple sequences. Presence or absence is noted by a horizontal line in the heatmap, and the relative copy number is noted by different tones of grey (darker in isolates with more copies, lighter in those with fewer). The dendrogram on the left of the heatmap highlights those isolates harbouring the same subset of genes.

A useful feature of Hound is its ability to populate iteratively the same assemblies to look for different genes, allowing the simultaneous detection of genes, calculation of their relative copy number, or re-analysis of prior data. Figure 3C shows two heatmaps illustrating the detection of different β−lactamases as well as nitroreductases, and efflux pumps—where mutations can cause resistance to multiple antibiotics [42]—in both farm and clinical isolates as well as any mutations found in their coding sequence and relative copy number.

## III. DISCUSSION

An increasingly problematic issue, particularly in the detection of antibiotic-resistance genes, is the lack of reproducibility [43, 44]. The choice of reference genomes is typically opaque to the user beyond the species, being notoriously difficult to underpin the exact assembly used. Along with the variety of existing pipelines that yield inconsistent results between laboratories [39], this problem led to the suggestion of standardised, ISO-certified pipelines [39]. Here we argue that they still fail to detect antibiotic resistance driven by the over-expression of enzymes like β−lactamases given their inability to account for gene over-expression caused by promoter mutations and increase in gene copy number. Now, beyond the detection of antibiotic resistance mutants, if we found different genetic signatures when comparing our assemblies to different references it is not unreasonable to think this will also be the case for other reference-mapped assemblies. Thus, using a standardised reference is unlikely to avoid this problem—exacerbated by the scarcity of tools to analyse *de novo* assemblies.

Hound is a step towards facilitating the analysis of these assemblies, not only by addressing a key limitation—gene fragmentation—but also by reducing the knowledge and technological burdens. The fact that we assembled and analysed more than 5,000 genomes on an 8-core, 16GB RAM laptop over the span of 9–10 days shows the potential for Hound to be implemented in larger and more powerful infrastructures for surveillance and diagnostic purposes. Now, Hound has some limitations. For example, it cannot report whether a gene of interest sits within a genomic island, but it can be used to detect whether genes associated to such islands are in the same contig as the gene of interest— complementing other bespoke analyses. Another limitation is that it currently only supports short-read Illumina sequencing data due to its availability during the development of this tool. However, given our use of SPAdes as backend assembler it is possible to add support for long-read nanopore and PacBio sequencing data in future releases.

The use of *de novo* assemblies means Hound is not only agnostic with respect to what genes can be monitored. It is also independent of the microbial species analysed—not possible with the use of reference genomes. With the majority of prokaryote diversity still being unknown and unsequenced [45, 46], we believe that Hound can be a useful tool to study non-model microorganisms that lack any reference—and help build them iteratively thanks to its contig-merging routine when sequencing costs in other platforms increase.

## IV. METHODS

### Genome sequencing and assembly

Genomic DNA libraries from isolate bacteria are prepared using the Nextera XT Library Prep Kit (Illumina, San Diego, USA) following the manufacturer’s protocol with the following modifications: Input DNA is increase 2-fold, and PCR elongation time is increased to 45 seconds. DNA quantification and library preparation are carried out on a Hamilton Microlab STAR automated liquid handling system (Hamilton Bonaduz AG, Switzerland), and the libraries sequenced on a Illumina NovaSeq 6000 (Illumina, San Diego, USA) using a 250 paired-end protocol by MicrobesNG. Reads are adapter-trimmed using Trimmomatic 0.30 [47] with a sliding window quality cutoff of Q15.

The flag `--preprocess reads.zip` processes the reads provider by MicrobesNG to create the directory structure required by Hound, with paired reads being stored in `$DIR/reads/`, and assemblies in `$DIR/assemblies/de_novo/` for reads assembled *de novo* or `$DIR/assemblies/reference-mapped/` for those mapped to a reference. This will depend on whether `--assembly --de-novo` or `--assembly --reference $REF_GENOME` have been passed. Hound then assembles *de novo* the resulting reads using SPAdes with the `--isolate` flag and *k*-mer size of 127, given the sequencing platform and protocol. For reference-mapped assemblies, Hound aligns the reads to a user-given reference using the Burrows-Wheeler Aligner and SAMtools with standard parameters. The coverage depth for all assemblies is then calculated with SAMtools if the flags `--coverage` and `--hk-genes $HK_GENES` are given, the baseline depth being the median coverage depth of all loci included in `$HK_GENES`, faster than computing the median coverage of the whole assembly to avoid any bias introduced by plasmid carriage. The relative copy number (RCN) is then calculated as the coverage depth of all loci in `$TARGET_GENES` divided by the baseline coverage [48].

### Indexing of assemblies

The resulting assemblies are indexed using `makeblastdb` from BLAST+, with flags `-parse_seqids` and `-dbtype nucl`, to facilitate the search of the sequences in file `$TARGET_GENES`. The search is run with `blastn`, which uses nucleotide sequences, or `tblastn` depending on whether the flag `--nucl` is passed to Hound. Without this flag, Hound assumes that the file `$TARGET_GENES` contains amino acid sequences and will therefore use `tblastn`.

### Multiple alignment and phylogeny

When the flag `--phylo` is passed, Hound will use muscle 3.80 to align the target gene sequences found in all assemblies given its accuracy and speed [49]. Penalties for the introduction and extension of gaps are pre-set with a value of `-9950.0` to avoid excessive fragmentation of the alignment. This alignment is then used by Hound to generate a phylogeny with PhyML [50] with seed `100100` for repeatability.

## Acknowledgements

Whole genome sequencing was performed by MicrobesNG. This work was funded by Medical Research Council grant MR/T005408/1 and Biotechnology and Biological Sciences Research Council Grant BB/X012670/1, and by grants MR/S004769/1, BB/T004592/1 and NE/N01961X/1 from the Antimicrobial Resistance Cross Council Initiative supported by the seven UK research councils, the Department of Health and Social Care and the National Institute for Health Research. It was further funded by grant 82459 from the Welsh Government Rural Communities -Rural Development Programme 2014-2020 supported by the European Union and the Welsh Government. N.S. was supported by a postgraduate scholarship from the University of Bristol.

## Code availability statement

A Python implementation of Hound can be found in https://gitlab.com/rc-reding/software/

## REFERENCES

1. McInerney, J. O., McNally, A. & O’connell, M. J. Why prokaryotes have pangenomes. Nat. Microbiol 2, 1–5 (2017).

2. Lan, R. & Reeves, P. R. Intraspecies variation in bacterial genomes: the need for a species genome concept. Trends Microbiol. 8, 396–401 (2000).

3. Venter, S. N., Palmer, M. & Steenkamp, E. T. Relevance of prokakryotic subspecies in the age of genomics. New Microbe and New Infect. 48 (2022).

4. Andam, C. P. Clonal yet different: understanding the causes of genomic heterogeneity in microbial species and impacts on public health. mSystems 4, e00097–19 (2019).

5. Tettelin, H., Masignani, V., Cieslewicz, M. J., et al. Genome analysis of multiple pathogenic isolates of Streptococcus agalactiae: implications for the microbial “pangenome”. Proc. Natl. Acad. Sci., USA 102, 13950–13955 (2005).

6. Tatusova, T., Ciufo, S., Fedorov, B., O’Neill, K. & Tolstoy, I. RefSeq microbial genomes database: new representation and annotation strategy. Nucleic Acids Res. 42, D553–D559 (2013).

7. Kaye, A. M. & Wasserman, W. W. The genome atlas: Navigating a new era of reference genomes. Trends Genet. 37, 807–818 (2021).

8. Clausen, P. T., Aarestrup, F. M. & Lund, O. Rapid and precise alignment of raw reads against redundant databases with KMA. BMC Bioinformatics 19, 1–8 (2018).

9. Guo, J., Bolduc, B., Zayer, A. A., et al. VirSorter2: a multi-classifier, expert-guided approach to detect diverse DNA and RNA viruses. Microbiome 9 (2021).

10. Rhie, A., McCarthy, S. A., Fedrigo, O., et al. Towards complete and error-free genome assemblies of all vertebrate species. Nature 592, 737–746 (2021).

11. Schneeberger, K., Ossowski, S., Ott, F., et al. Reference-guided assembly of four diverse Arabidopsis thaliana genomes. Proc. Natl. Acad. Sci., USA 108, 10249–10254 (2011).

12. Formenti, G., Theissinger, K., Fernandes, C., et al. The era of reference genomes in conservation genomics. Trends Ecol. Evol. 37, 197–202 (2022).

13. Didelot, X., Bowden, R., Wilson, D. J., Peto, T. E. & Crook, D. W. Transforming clinical microbiology with bacterial genome sequencing. Nat. Rev. Genet. 13, 601–612 (2012).

14. Pankhurst, L. J., del Ojo Elias, C., Votintseva, A. A., et al. Rapid, comprehensive, and affordable mycobacterial diagnosis with whole-genome sequencing: a prospective study. Lancet Respir. Med. 4, 49–58 (2016).

15. Hendriksen, R. S., Bortolaia, V., Tate, H., Tyson, G. H., Aarestrup, F. M. & McDermott, P. F. Using genomics to track global antimicrobial resistance. Front. Public Health 7, 242 (2019).

16. Sichtig, H., Minogue, T., Yan, Y., et al. FDA-ARGOS is a database with public qualitycontrolled reference genomes for diagnostic use and regulatory science. Nat. Commun. 10, 3313 (2019).

17. GLASS whole-genome sequencing for surveillance of antimicrobial resistance World Health Organization (World Health Organization, 2020). isbn: 978-92-4-001100-7.

18. Walker, T. M., Kohl, T. A., Omar, S. V., et al. Whole-genome sequencing for prediction of Mycobacterium tuberculosis drug susceptibility and resistance: a retrospective cohort study. Lancet Infect. Dis. 15, 1193–1202 (2015).

19. Su, M., Satola, S. W. & Read, T. D. Genome-Based Prediction of Bacterial Antibiotic Resistance. J. Clin. Microbiol. 57, e01405–18 (2019).

20. Valiente-Mullor, C., Beamud, B., Ansari, I., et al. One is not enough: on the effects of reference genome for the mapping and subsequent analyses of short-reads. PLoS Comput. Biol. 17, e1008678 (2021).

21. Mukherjee, S., Seshadri, R., Varghese, N. J., et al. 1,003 reference genomes of bacterial and archaeal isolates expand coverage of the tree of life. Nat. biotechnol. 35, 676–683 (2017).

22. Zankari, E., Hasman, H., Cosentino, S., et al. Identification of acquired antimicrobial resistance genes. J. Antimicrob. Chemother. 67, 2640–2644 (2012).

23. De Man, T. J. & Limbago, B. M. SSTAR, a stand-alone easy-to-use antimicrobial resistance gene predictor. mSphere 1, 10–1128 (2016).

24. Gupta, S. K., Padmanabhan, B. R., Diene, S. M., et al. ARG-ANNOT, a new bioinformatic tool to discover antibiotic resistance genes in bacterial genomes. Antimicrob. Agents Chemother. 58, 212–220 (2014).

25. Davis, J. J., Boisvert, S., Brettin, T., et al. Antimicrobial resistance prediction in PATRIC and RAST. Sci. Rep. 6, 27930 (2016).

26. Seemann, T. ABRIcate eprint: https://github.com/tseemann/abricate.

27. Fuentes-Pardo, A. P. & Ruzzante, D. E. Whole-genome sequencing approaches for conservation biology: Advantages, limitations and practical recommendations. Mol. Ecol. 26, 5369–5406 (2017).

28. Bailey, J. K., Pinyon, J. L., Anantham, S. & Hall, R. M. Distribution of the blaTEM gene and blaTEM-containing transposons in commensal Escherichia coli. J. Antimicrob. Chemother. 66, 745–751 (2011).

29. Davies, T. J., Stoesser, N., Sheppard, A. E., et al. Reconciling the potentially irreconcilable? Genotypic and phenotypic amoxicillin-clavulanate resistance in Escherichia coli. Antimicrob. Agents Chemother. 64, 10–1128 (2020).

30. Ruffles, T. J., Goyal, V., Marchant, J. M., et al. Duration of amoxicillin-clavulanate for protracted bacterial bronchitis in children (DACS): a multi-centre, double blind, randomised controlled trial. Lancet Respir. Med. 9, 1121–1129 (2021).

31. KuKanich, K., Lubbers, B. & Salgado, B. Amoxicillin and amoxicillin-clavulanate resistance in urinary Escherichia coli antibiograms of cats and dogs from the Midwestern United States. J. Vet. Intern. Med. 34, 227–231 (2020).

32. Hansen, K. H., Andreasen, M. R., Pedersen, M. S., Westh, H., Jelsbak, L. & Schønning, K. Resistance to piperacillin/tazobactam in Escherichia coli resulting from extensive IS 26-associated gene amplification of blaTEM-1. J. Antimicrob. Chemother. 74, 3179–3183 (2019).

33. Bankevich, A., Nurk, S., Antipov, D., et al. SPAdes: a new genome assembly algorithm and its applications to single-cell sequencing. J. Comput. Biol. 19, 455–477 (2012).

34. Li, H. & Durbin, R. Fast and accurate short read alignment with Burrows–Wheeler transform. Bioinformatics 25, 1754–1760 (2009).

35. Li, H., Handsaker, B., Wysoker, A., et al. The sequence alignment/map format and SAMtools. Bioinformatics 25, 2078–2079 (2009).

36. Camacho, C., Coulouris, G., Avagyan, V., et al. BLAST+: architecture and applications. BMC Bioinformatics 10, 1–9 (2009).

37. Wood, D. E., Lu, J. & Langmead, B. Improved metagenomic analysis with Kraken 2. Genome biol. 20, 1–13 (2019).

38. Delcher, A. L., Phillippy, A., Carlton, J. & Salzberg, S. L. Fast algorithms for large-scale genome alignment and comparison. Nucleic Acids Res. 30, 2478–2483 (2002).

39. Sherry, N. L., Horan, K. A., Ballard, S. A., et al. An ISO-certified genomics workflow for identification and surveillance of antimicrobial resistance. Nat. Commun. 14, 60 (2023).

40. Mounsey, O., Marchetti, L., Parada, J., et al. Genomic epidemiology of thirdgeneration cephalosporin-resistant Escherichia coli from Argentinian pig and dairy farms reveals animal-specific patterns of co-resistance and resistance mechanisms. bioRxiv. 10.1101/2023.06.15.545115 (2023).

41. San Millan, A., Escudero, J. A., Gifford, D. R., Mazel, D. & MacLean, R. C. Multicopy plasmids potentiate the evolution of antibiotic resistance in bacteria. Nat. Ecol. Evol. 1, 0010 (2016).

42. Blair, J. M., Bavro, V. N., Ricci, V., et al. AcrB drug-binding pocket substitution confers clinically relevant resistance and altered substrate specificity. Proc. Natl. Acad. Sci. U.S.A. 112, 3511–3516 (2015).

43. Doyle, R. M., O’sullivan, D. M., Aller, S. D., et al. Discordant bioinformatic predictions of antimicrobial resistance from whole-genome sequencing data of bacterial isolates: an inter-laboratory study. Microbial genomics 6, e000335 (2020).

44. Coolen, J. P., Jamin, C., Savelkoul, P. H., et al. Centre-specific bacterial pathogen typing affects infection-control decision making. Microb. Genom. 7, 000612 (2021).

45. Rappé, M. S. & Giovannoni, S. J. The uncultured microbial majority. Annu. Rev. Microbiol. 57, 369–394 (2003).

46. Locey, K. J. & Lennon, J. T. Scaling laws predict global microbial diversity. Proc. Natl. Acad. Sci. U.S.A. 113, 5970–5975 (2016).

47. Bolger, A. M., Lohse, M. & Usadel, B. Trimmomatic: a flexible trimmer for Illumina sequence data. Bioinformatics 30, 2114–2120 (2014).

48. Reding, C., Catalán, P., Jansen, G., et al. The Antibiotic Dosage of Fastest Resistance Evolution: gene amplifications underpinning the inverted-U. Mol. Biol. Evol. 9, 3847–3863. issn: 0737-4038 (Mar. 2021).

49. Edgar, R. C. MUSCLE: a multiple sequence alignment method with reduced time and space complexity. BMC Bioinformatics 5, 1–19 (2004).

50. Guindon, S. & Gascuel, O. A simple, fast, and accurate algorithm to estimate large phylogenies by maximum likelihood. Syst. Biol. 52, 696–704 (2003).

